# Bulk and Single-nucleus Transcriptomics Highlight Intra-telencephalic and Somatostatin Neurons in Alzheimer’s Disease

**DOI:** 10.1101/2022.01.12.476076

**Authors:** Micaela E Consens, Yuxiao Chen, Vilas Menon, Yanling Wang, Julie A Schneider, Philip L De Jager, David A Bennett, Shreejoy J Tripathy, Daniel Felsky

**Affiliations:** The Krembil Centre for Neuroinformatics (KCNI), Centre for Addiction and Mental Health (CAMH), Toronto, ON, CA; The Rush Alzheimer’s Disease Center, Rush University, Chicago, Il, USA; The Centre for Translational and Computational Neuroimmunology, Columbia University Medical Center, New York, NY, USA; Department of Psychiatry, University of Toronto, Toronto, ON, CA; Institute of Medical Science, University of Toronto, Toronto, ON, CA; Dalla Lana School of Public Health, University of Toronto, Toronto ON, CA; Department of Physiology, University of Toronto, Toronto, ON, CA

**Keywords:** Cell type proportions, Alzheimer’s disease, somatostatin, RNA sequencing, postmortem brain, mega-analysis

## Abstract

**Background:** Cortical neuron loss is a pathological hallmark of late-onset Alzheimer’s disease (AD). However, it remains unclear which neuronal subtypes are most vulnerable to degeneration and contribute most to cognitive decline.

**Methods:** We analyzed postmortem bulk brain RNA-sequencing (RNAseq) data collected from three studies of aging and AD comprising six neocortical regions (704 individuals; 1037 samples). We estimated relative cell type proportions from each brain sample using neuronal subclass-specific marker genes derived from ultra-high depth single-nucleus RNAseq data (snRNAseq). We associated cell type proportions with AD across all samples using mixed-effects mega-analyses. Bulk tissue analyses were complemented by analyses of three AD snRNAseq datasets using the same cell type definitions and diagnostic criteria (51 individuals). Lastly, we identified cell subtype associations with specific neuropathologies, cognitive decline, and residual cognition.

**Results:** In our mega-analyses, we identified the strongest associations of AD with fewer somatostatin (SST) inhibitory neurons (β=−0.48, *p*_bonf_=8.98×10^−9^) and intra-telencephalic (IT) excitatory neurons (β=−0.45, *p*_bonf_ =4.32×10^−7^). snRNAseq-based cell type proportion analyses especially supported the association of SST neurons. Analyses of cell type proportions with specific AD-related phenotypes in ROS/MAP consistently implicated fewer SST neurons with greater brain-wide postmortem tau and beta amyloid (β=−0.155, *p*_FDR_=3.1×10^−4^) deposition, as well as more severe cognitive decline prior to death (β=0.309, *p*_FDR_=3.9×10^−6^). Greater IT neuron proportions were associated strongly with improved cognition (β=0.173, *p*_FDR_=8.3×10^−5^) and residual cognition (β=0.175, *p*_FDR_=1.2×10^−5^), but not canonical AD neuropathology.

**Conclusions:** Proportionally fewer SST and IT neurons were significantly associated with AD diagnosis across multiple studies and cortical regions. These findings support seminal work implicating somatostatin and pyramidal neurons in the pathogenesis of AD and improves our current understanding of neuronal vulnerability in AD.

## Background

Late-onset Alzheimer’s disease (AD) is a neurodegenerative disease characterized by the gradual accumulation of specific neuropathologies, including beta amyloid and hyperphosphorylated tau proteins, followed by widespread brain atrophy and cognitive decline [1–3]. While these pathological hallmarks of AD are well established, a lack of clarity over which specific brain cell types are lost over the course of neurodegeneration and cognitive decline remains.

Recent advances in single-cell and cell type-specific gene expression profiling has revolutionized our knowledge of cell-type specific changes in neuropsychiatric disease [4–8]. By combining these datasets with bulk tissue gene expression data sampled from large numbers of well-characterized individuals, cellular deconvolution analyses have revealed important differences in AD, including fewer neurons and more astrocytes [9–11]. However, the majority of these analyses have focused on cellular differences at the level of broad cell types and comparatively less focus has been placed on resolving cellular differences among finer-resolution cell types such as subtypes of neurons [12]. While emerging AD case/control snRNAseq datasets offer an exciting opportunity to better resolve such cellular differences [12–15], technical constraints have limited the size of such datasets in terms of total numbers of cells and individuals sampled [9], making it difficult to determine robust cellular differences in a disorder as heterogeneous as AD.

Here we performed a comprehensive analysis of brain bulk- and single-nucleus RNAseq datasets to quantify changes in cell-type proportions in AD. We quantified excitatory and inhibitory neuronal subpopulations and non-neuronal cell types by estimating relative cell-type proportions across three studies and six neocortical brain regions. We corroborated our bulk tissue-based findings by directly estimating cell-type proportions in three snRNAseq datasets collected from AD cases and controls. Finally, we explored how cell-type proportion differences relate to specific age-related neuropathologies, longitudinal cognitive decline, and an established measure of cognitive resilience. Together, our findings suggest a robust and specific loss of excitatory intra-telencephalic neurons and inhibitory somatostatin-expressing interneurons in AD.

## Methods

### Studies used for bulk tissue RNAseq analyses

Postmortem bulk-brain RNAseq data was processed from 1 373 different individuals across three independent studies from the Accelerating Medicines Partnership - Alzheimer’s Disease (AMP-AD) consortium (summarized in **Table 1**), encompassing six brain regions:

**Table 1.**
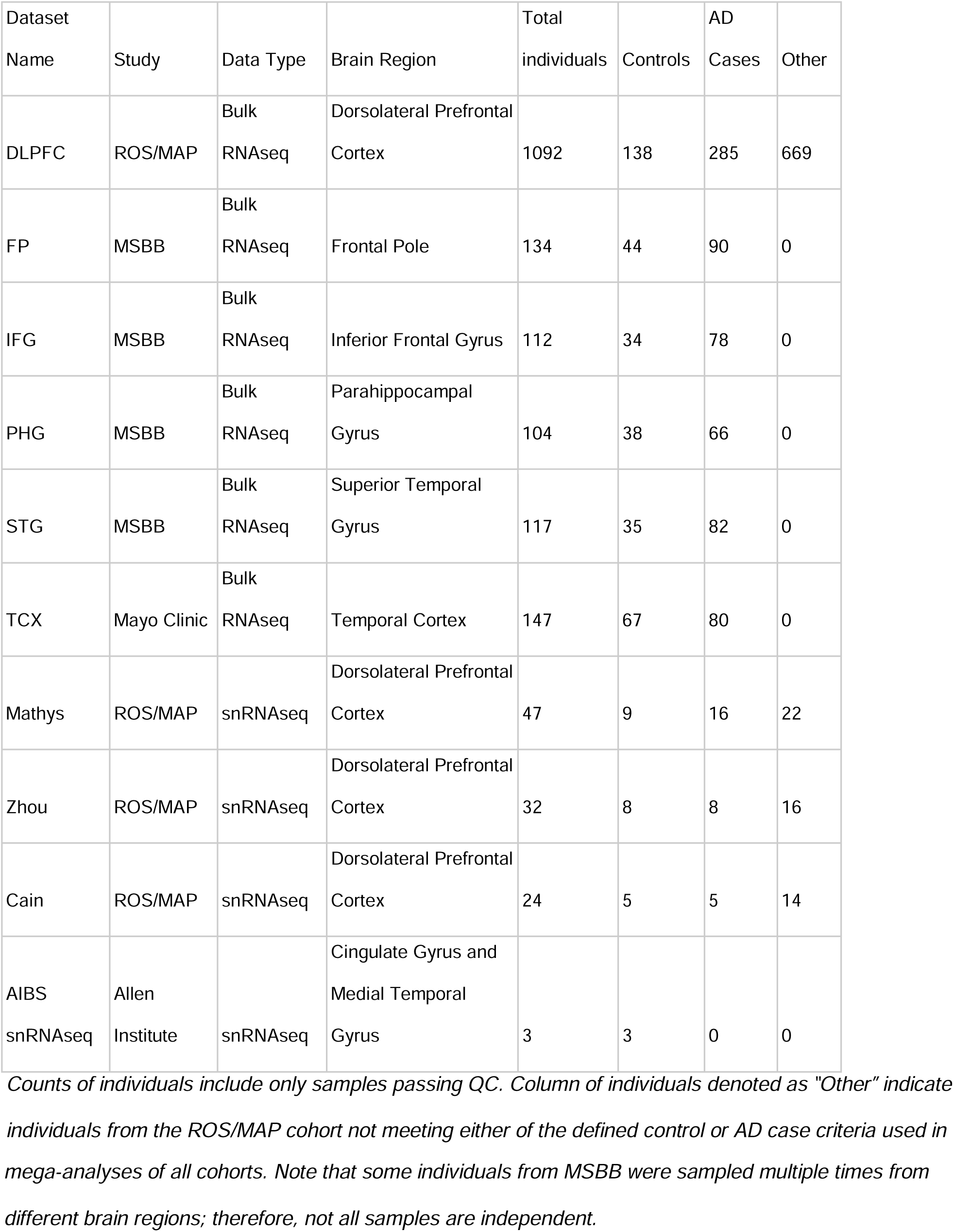
Summary of RNAseq datasets used in this study.

| Dataset Name | Study | Data Type | Brain Region | Total individuals | Controls | AD Cases | Other |
| --- | --- | --- | --- | --- | --- | --- | --- |
| DLPFC | ROS/MAP | Bulk RNAseq | Dorsolateral Prefrontal Cortex | 1092 | 138 | 285 | 669 |
| FP | MSBB | Bulk RNAseq | Frontal Pole | 134 | 44 | 90 | 0 |
| IFG | MSBB | Bulk RNAseq | Inferior Frontal Gyrus | 112 | 34 | 78 | 0 |
| PHG | MSBB | Bulk RNAseq | Parahippocampal Gyrus | 104 | 38 | 66 | 0 |
| STG | MSBB | Bulk RNAseq | Superior Temporal Gyrus | 117 | 35 | 82 | 0 |
| TCX | Mayo Clinic | Bulk RNAseq | Temporal Cortex | 147 | 67 | 80 | 0 |
| Mathys | ROS/MAP | snRNAseq | Dorsolateral Prefrontal Cortex | 47 | 9 | 16 | 22 |
| Zhou | ROS/MAP | snRNAseq | Dorsolateral Prefrontal Cortex | 32 | 8 | 8 | 16 |
| Cain | ROS/MAP | snRNAseq | Dorsolateral Prefrontal Cortex | 24 | 5 | 5 | 14 |
| AIBS snRNAseq | Allen Institute | snRNAseq | Cingulate Gyrus and Medial Temporal Gyrus | 3 | 3 | 0 | 0 |

Counts of individuals include only samples passing QC. Column of individuals denoted as “Other” indicate individuals from the ROS/MAP cohort not meeting either of the defined control or AD case criteria used in
*mega-analyses of all cohorts. Note that some individuals from MSBB were sampled multiple times from* *different brain regions; therefore, not all samples are independent.*

1. The Religious Orders Study and Rush Memory and Aging Project (herein ROS/MAP) cohort provided bulk RNAseq data for dorsolateral prefrontal cortex (DLPFC) from 1092 individuals. The mean age at death was 89.6 (SD=6.6).
2. The Mayo Clinic study (herein Mayo) provided temporal cortex (TCX) samples from 147 individuals. The mean age at death was 82.6 (SD=8.0).
3. The Mount Sinai Brain Bank study (herein MSBB) provided samples from the same individuals across multiple brain regions. The mean age at death was 83.3 (SD=7.4). 134 individuals had bulk-tissue RNAseq data sampled from Frontal Pole (FP), Brodmann area 10; 112 from Inferior Frontal Gyrus (IFG), Brodmann area 44 (BA44); 104 from Parahippocampal Gyrus (PHG), Brodmann area 36; and 117 from Superior Temporal Gyrus (STG), Brodmann area 22 (BA22).

### Tissue preparation and bulk tissue RNA sequencing

Details pertaining to the handling and processing of postmortem samples in ROS/MAP [16], Mayo [17], and MSBB [18] have been previously published [19]. RNA sequencing procedures differed between studies:

1. For ROS/MAP, RNA sequencing on DLPFC tissue was carried out in 13 batches within three distinct library preparation and sequencing pipelines (see **Supplementary Methods**). Sequencing was carried out using the Illumina HiSeq (pipeline #1: 50M 101bp paired end reads) and NovaSeq6000 (pipeline #2: 30M 100bp paired end; pipeline #3: 40-50M 150bp paired end reads). Full details on RNA extraction and sequencing are available on the Synapse AMP-AD Knowledge Portal (syn3219045).
2. For Mayo, sequencing was carried out on the Illumina HiSeq 2000 (101bp paired end reads). Details available on the AMP-AD Knowledge Portal (syn5550404)
3. For MSBB, sequencing was carried out on the Illumina HiSeq 2500 (100bp single end reads).

Details available on the AMP-AD Knowledge Portal (syn3159438)

### Processing of bulk tissue RNAseq datasets

Bulk-tissue based RNA-seq read counts from all three studies underwent uniform quality control (QC) and filtering protocols, using the same approach as described previously [20]. Briefly, genes with a median expected count less than or equal to 15 were removed and multidimensional scaling was performed. Subjects were deemed outliers and removed if they differed from the sample median of any of the first 5 latent components by more than 3 interquartile ranges. Gene counts were log2 transformed with an offset of 0.5, to coerce any log2(expected count) value differing from the sample median by 3 interquartile ranges to its nearest most extreme value within that range. After sample- and gene-level filtering, the log2(expected counts) were transformed back to the expected count scale. Trimmed mean of m-values (TMM) normalization (using edgeR calcNormFactors) and mean-variance derived observational-level weights were calculated. Variance related to technical factors, including sequencing batch, percent of mapped bases, percent usable bases, RNA integrity number (RIN), and postmortem interval, were removed using the removeBatchEffect function in limma [21].

### Consensus Definition of Alzheimer’s Disease for Mega-Analysis

We applied a harmonized definition of AD case/control diagnosis as defined previously in the same studies [19]. This definition combined clinical and postmortem neuropathological data for categorization, where controls were defined as individuals with a low burden of plaques and tangles, as well as no evidence of cognitive impairment (if available). To define AD case status, Braak stage, global cognitive status at time of death, and CERAD (Consortium to Establish a Registry for Alzheimer’s Disease) scores were used in ROS/MAP, with Clinical Dementia Rating (CDR) scores being used instead of global cognitive status in MSBB. For the Mayo dataset, only neuropathological criteria were used to define case/control status, with details previously published [22]. In total, 704 individuals across the three studies met the established AD case or control criteria.

### Cognitive and Neuropathological measures in ROS/MAP

All subjects in ROS/MAP were administered 17 cognitive tests annually spanning five cognitive domains. Raw scores for tests within each domain were z-scored (using the mean and standard deviation of the entire ROS/MAP combined study population at baseline) and averaged to form the composites. The list of individual cognitive tasks and their corresponding domains has been published [23]. Prior to autopsy, the average postmortem interval was 9.3 hours (SD=8.1). All brains were examined by a board-certified neuropathologist blinded to clinical data. We analyzed 11 neuropathologies measured brain-wide: total Aβ peptides, neuritic and diffuse plaques, paired helical filament tau protein, neurofibrillary tangles, Braak stage (tau), gross cerebral infarcts, cerebral atherosclerosis, degree of alpha-synucleinopathy, TDP43 proteinopathy, and hippocampal sclerosis. Detailed descriptions of all neuropathological variables have been previously published [23].

### Single-nucleus RNAseq datasets

In total, we used expression data from four human cortical single-nucleus RNA sequencing (snRNAseq) datasets for this study. First, we used an ultra high-depth SMART-seq based snRNAseq dataset from the human neocortex provided by the Allen Institute for Brain Sciences (AIBS) [24] to define our reference cell type taxonomy and derive cell type specific marker genes (see **Supplemental Methods**). We used all nuclei sampled from the cingulate gyrus (5,939 nuclei) and medial temporal cortex (15,519), as these correspond most closely with the bulk expression samples described above. Given that nuclei from non-neuronal cell types were relatively undersampled in this dataset, we supplemented this dataset with 2,620 nuclei corresponding to non-neurons sampled from other cortical regions, including visual, auditory, somatosensory and motor cortex (502, 742, 595, and 781 nuclei respectively). We further used three AD case/control snRNAseq datasets collected from subjects sampled from the ROS/MAP cohort [12-14]. Cells from each of the three ROS/MAP snRNAseq datasets were bioinformatically mapped onto the AIBS snRNAseq dataset (see **Supplemental Methods**).

### Estimation of relative cell type proportions from bulk RNAseq samples

Relative cell type proportions were estimated with the MarkerGeneProfile (MGP) R package, as described previously [4,5], using our derived cell type-specific marker genes with default parameters. The output of the mgpEstimate function was taken as the relative cell-type proportion estimates (rCTPs), providing an indirect measure of cell type abundance in each sample. To ensure consistency in rCTP definitions across individual bulk datasets, rCTPs were estimated using only cell type-specific marker genes passing QC in each of the six bulk-tissue datasets. rCTPs were converted to standardized z-scores within each dataset prior to downstream analysis.

### Estimation of snRNAseq-derived cell type proportions

Cell type proportions from snRNAseq datasets (snCTPs) were directly estimated from snRNAseq datasets by counting nuclei annotated to each cell type and normalizing by the total count of all QC-passing nuclei per individual subject. We note that such calculations were only performed on nuclei passing quality control and also met our mapping criteria to our reference cell type taxonomy. Direct comparisons between bulk and snRNAseq derived cell type proportions for subjects from the ROS/MAP cohort were performed by identifying subjects in common between both datasets and correlating rCTPs with snCTPs values across subjects.

### Statistical Analyses

#### Mega-analysis of Bulk RNA-seq cell type proportions with AD

The lme4 package in R [25] was used to perform a set of mega-analyses, one per cell type, across all bulk RNAseq datasets. Linear mixed effect models were fitted as follows, for each cell type (i), including a random effect of sample to account for correlation structure between brain samples taken from multiple regions of the same individuals in the MSBB study:

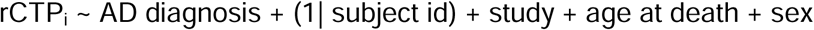

Adjustment of two-sided *p*-values to account for multiple cell types was performed. The Bonferroni method and false discovery-rate (FDR) method [26] were applied separately to identify highly stringent and more relaxed thresholds for statistical significance. Corrected *p*-values are labelled specifically within results (i.e. *p*_Bonf_, *p*_FDR_).

#### Analysis of single-nucleus CTPs in AD and controls

For snCTPs, AD association analyses were performed as for rCTPs, with an additional covariate added for PMI (for bulk analyses, variation in gene expression due to PMI was removed during the preprocessing phase) and using a linear model due to the limited overlap in individuals sampled between snRNAseq datasets.

#### Association of bulk tissue rCTPs with neuropathology, cognitive decline, and residual cognition in ROS/MAP

In ROS/MAP we performed detailed analyses associating each rCTP to measures of individual brain pathologies, global cognitive decline, and cognitive status proximal to death. We specifically utilized the full set of individuals in ROS/MAP with bulk expression and other measures available, as opposed to the more limited set of individuals in our cross-cohort mega-analysis of AD case/control criteria described above. “Residual cognition” was calculated per individual as in White et al.[27]; the measure corresponds to residuals of a linear model regressing global cognition proximal to death on observed neuropathological variables and demographic factors. Associations between rCTPs for each cell type and cognitive and neuropathological phenotypes were assessed using linear models covarying for age at death, sex and pmi. For models of cognitive outcomes, we covaried for sex, educational attainment, and age at time of baseline study assessment. Correction for multiple testing across all cell types and outcomes (19*14=266 tests) was performed using the FDR method. To estimate variance explained (R^2^) by rCTPs over and above covariates and measured neuropathologies, we built a series of nested linear models and compared them using likelihood ratio tests. To improve the generalizability of our estimates, models were bootstrapped (100 iterations) using the .632+ method [28].

#### Causal mediation modelling of IT and SST rCTPs, AD neuropathology, and cognitive performance in ROS/MAP

The R mediation package (v4.5.0) was used for causal mediation modelling. To model pathological burden, we used an established measure of global postmortem AD neuropathology: the mean of brain-wide diffuse plaques, neuritic plaques, and neurofibrillary tangles. Four models were tested with SST and IT rCTPs as either predictor or mediator, AD pathology the other role and global cognitive performance at last study visit was always the outcome measure. To estimate confidence intervals for average indirect, direct, and total effects, 1000 Monte Carlo draws were used for nonparametric bootstrapping.

## Results

### Deriving cell type-enriched marker genes for the major neuron subclasses of the human neocortex

To build a high-quality foundation for investigating cell subtype proportions - focusing on subclasses of neocortical neurons - in AD, we first established representative marker gene sets using ultra-high depth single-nucleus data RNA sequencing (snRNAseq) data from multiple regions of human cortex collected by the Allen Institute for Brain Sciences [24]. In these datasets, nuclei were annotated to an established cell type reference taxonomy where transcriptomically-defined cell clusters are linked to orthologous multi-modal taxonomies established in other species and neocortical regions [29–31]. This annotation enables the inference of additional aspects of cellular identity for these human snRNAseq-based cell clusters that include axonal projection patterns and cell morphology [24].

We used these snRNAseq reference data to derive cell type-specific “marker genes’’ (illustrated in **Supplementary Table 1**), focusing our analyses primarily on the subclass cell type resolution. This taxonomic grouping serves as an intermediate resolution (e.g. somatostatin-expressing GABAergic interneurons) between more coarse-grained (e.g. inhibitory neurons) and fine-grained cell taxonomic divisions (e.g. Martinotti neurons). A key benefit of these markers is their specificity and good cross-dataset replicability, including in snRNAseq datasets specific to aging and AD samples (**Supplementary Figures 1 and 2**).

### Bulk tissue analysis implicates fewer inhibitory and excitatory neurons in AD, with most robust associations with SST interneurons and IT pyramidal cells

We first sought to understand how the abundance of specific cell types are different in brains of individuals with a pathologically-confirmed AD diagnosis compared to controls. We estimated the relative cell type proportions (rCTPs) of each post-mortem bulk tissue RNAseq sample across all six bulk expression datasets separately using the Marker Gene Profile (MGP) method [4,5] and our novel cell type-enriched marker gene sets described above. We then performed mega-analysis for AD case/control status with rCTPs across each of the major cell subclasses and all six datasets using a linear mixed effects model. In aggregate, we found lower rCTPs in most GABAergic subclasses, mostly fewer but some greater rCTPs among glutamatergic subclasses, and higher rCTPs for most non-neuronal subclasses (**Figure 1A**).

**Figure 1:**
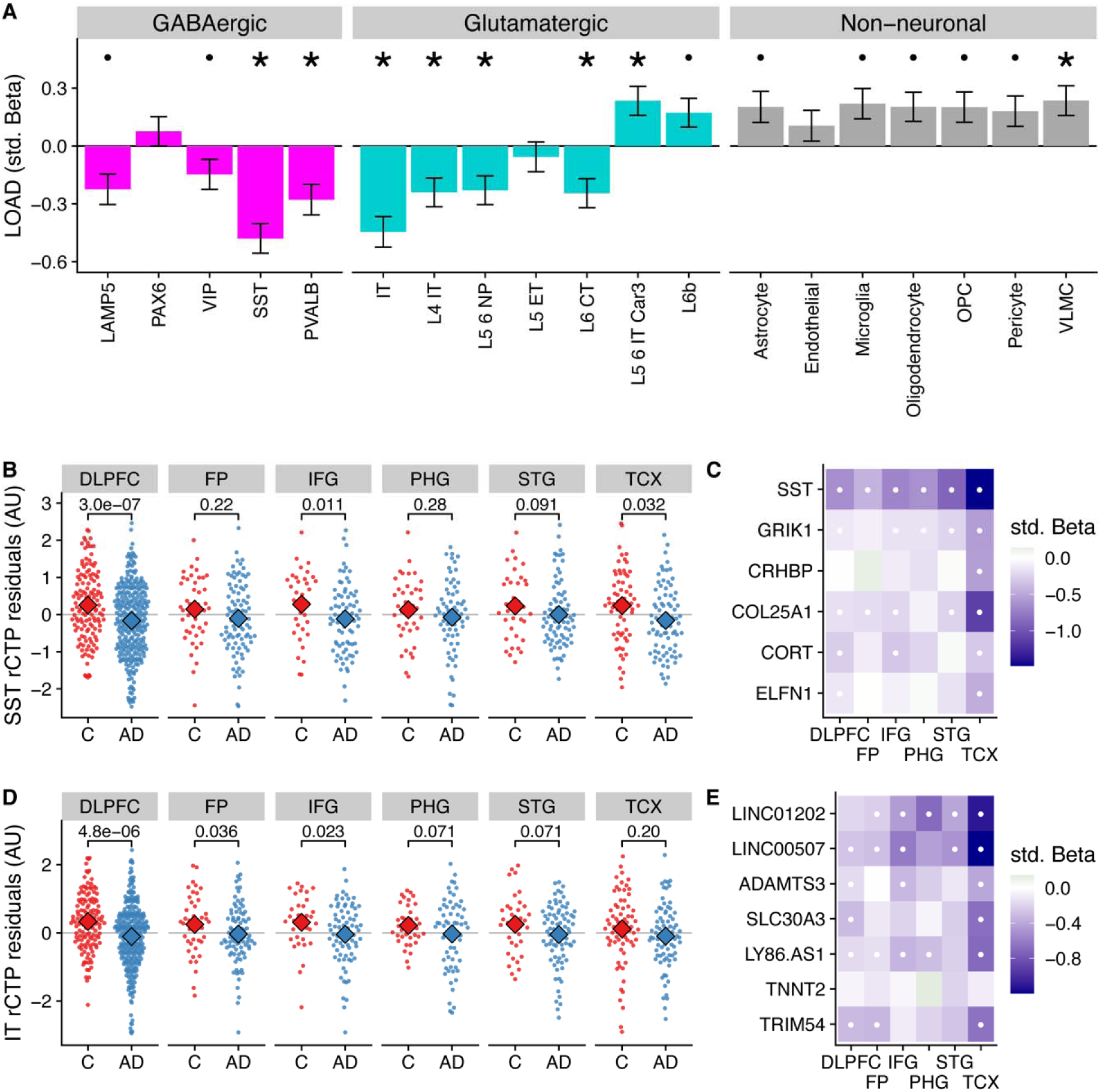
Differences in relative cell type proportions of neuronal and non-neuronal subclasses in Alzheimer’s disease. A) Consensus associations of AD status and cell type-specific relative cell type proportions (rCTPs) across six bulk RNAseq datasets. Y-axis shows standardized beta coefficients estimated using a mixed effects model to pool associations across datasets. Positive (negative) standardized beta coefficients indicate an increase (decrease) in the cell type-specific rCTP in AD. Error bars indicate one standard deviation. Asterisks (dots) above each cell type indicate two-sided p_bonf_ < 0.05 (or FDR < 0.1). B) Comparisons between rCTPs between controls and AD cases in each of the six bulk gene expression datasets, ROS/MAP, sampling the dorsolateral prefrontal cortex (DLPFC), MSSB, sampling the Frontal Pole (FP), Inferior Frontal Gyrus (IFG), Parahippocampal Gyrus (PHG), and Superior Temporal Gyrus (STG), and the Mayo cohort, sampling the Temporal Cortex (TCX). Y-axis shows estimates of rCTPs for somatostatin (SST) interneurons from individual post-mortem samples (each dot reflects one individual), after covarying for age and sex. Numbers show p-values from a statistical model comparing residualized rCTPs between controls and AD cases, uncorrected for multiple comparisons across datasets and cell types. Subjects with outlier values of rCTPs not shown. C) Heatmaps showing AD case/control associations for marker genes for SST inhibitory cells. White dots indicate specific associations where FDR < 0.1. D, E) Same as B, C for intra-telencephalic-projecting (IT) excitatory pyramidal cells (IT cells).

Specifically, at a stringent Bonferroni correction threshold of *p*_bonf_ < 0.05, our analysis identified lower rCTPs for SST (β=−0.48, *p*_bonf_=8.98*10^−9^) and PVALB (β=−0.28, *p*_bonf_=0.0072) GABAergic interneurons, as well as IT (β=−0.45, *p*_bonf_ =4.32*10^−7^), L4 IT (β = -0.24, p_bonf_ = 0.023), L5 6 NP (β=−0.23, *p*_bonf_=0.039) and L6 CT (β=−0.25, *p*_bonf_=0.023) glutamatergic neurons in AD. At the same threshold, we observed greater rCTPs for L5 6 IT Car3 glutamatergic neurons (β=0.023, *p*_bonf_=0.034) and VLMC cells (β=0.24, *p*_bonf_=0.041). At a less stringent threshold (FDR<0.1), we also observed lower rCTPs for LAMP5 and VIP GABAergic interneurons, and greater rCTPs for L6b glutamatergic cells and most non-neuronal cells, except endothelial cells. One important caveat of these analyses is the focus on relative proportions, which are not absolute cell counts [4,5]; therefore, potentially paradoxical reported differences in some rCTPs here, such as greater L5 6 IT Car3 and L6b glutamatergic cell rCTPs in AD, may not necessarily indicate that these cell types are actually increasing in their absolute cell counts. These results are consistent with prior observations that AD is characterized by relatively fewer neuronal cells and corresponding relatively more non-neuronal cells [32].

The strongest AD-associated cell type in mega-analysis was the somatostatin (SST) interneuron (β=−0.48, *p*_bonf_=8.98*10^−9^); in each individual dataset, SST rCTPs were lower in AD cases relative to controls (**Figure 1B**), though the differences were not significant in all regions. Our findings mirror those of Cain et al. [12] highlighting SST interneurons as particularly associated with AD phenotypes among ROS/MAP participants and further generalizes this finding to additional studies and brain regions. Importantly, in addition to the *SST* gene mRNA transcript itself, we also observed lower mRNA expression of other SST interneuron marker genes, including *GRIK1* and *COL25A1* across most brain regions (**Figure 1C**). Moreover, the SST rCTP signal is robust, albeit attenuated, to the removal of the SST mRNA as a marker gene (β=−0.39, *p*_bonf_=6.02*10^−6^), suggesting the relevance of fewer SST-expressing neurons rather than a lower expression of the SST gene specifically.

Among excitatory cell types, rCTPs for intratelencephalic-projecting (IT) pyramidal cells showed the greatest difference between AD and controls (β=−0.45, *p*_bonf_=4.32*10^−7^). Like SST rCTPs, proportionally fewer IT neurons were observed in AD cases relative to controls in each of the six bulk expression datasets (**Figure 1D**), albeit not significantly in all regions. The IT cell subclass includes both superficial layer pyramidal cells, such as CUX2-positive cells, as well as more infragranular cells, including RORB- and THEMIS-positive neurons [24]. As expected, we observed lower expression in many of the individual IT cell marker genes across each of the datasets in AD, including *LINC00507* and *LINC01202* (**Figure 1E**).

### Single-nucleus analysis suggests selective vulnerability of specific inhibitory subclasses in AD, including LAMP5 and SST interneurons, but not IT-projecting pyramidal cells

To complement the indirect estimates of rCTPs from bulk tissue, we accessed three AD case/control snRNAseq datasets sampled from subsets of participants from the ROS/MAP cohort (**Table 1**) [12–14]. We first harmonized cell type annotations from these snRNAseq datasets by mapping cells to the same Allen Institute-based human cortical cell type reference taxonomy used in our rCTP analyses, finding good qualitative agreement between the annotated cell type identities provided within the original publications and those following QC and re-mapping (**Supplemental Figure 3**). We then estimated single-nucleus CTPs (snCTPs) per post-mortem sample by counting nuclei annotated to each cell type, as a percentage of the total nuclei sampled in each subject. Correlations between cell type-specific snCTPs and bulk-tissue derived rCTPs in subjects with overlapping data types were modest, but overall positive (**Supplementary Figure 4**), in agreement with published deconvolution efforts validated using immuno-histochemistry [32].

In line with our bulk tissue rCTP analysis, a mega-analysis across the three snRNAseq datasets indicated that AD samples showed lower snCTPs in most inhibitory subclasses, both higher and lower snCTPs among excitatory subclasses, and greater snCTPs for most non-neuronal subclasses (**Figure 2A**). At Bonferroni-corrected *p*_bonf_<0.05, we found only LAMP5 GABAergic interneurons to be lower in AD compared to controls (β=−0.95, *p*_bonf_=0.011). At a less stringent threshold (FDR<0.1), we also observed lower snCTPs for PAX6 (β=−0.62, FDR=0.093) and SST (β=−0.74, FDR=0.093) interneurons (**Figure 2B**). We note that we did not see strong evidence for lower expression of SST mRNA among SST-annotated nuclei in AD after covarying for SST cell proportion differences (β=−0.43, p = 0.47; **Supplementary Figure 5**), providing additional evidence for SST cell-specific vulnerability highlighted by bulk tissue analysis. In contrast to our bulk tissue results, we did not find any effects for (IT) pyramidal cells (**Figure 2C**). To assess the overall consistency between our bulk tissue rCTP and single-nucleus approaches, we correlated standardized effect coefficients for each cell type between analyses (**Figure 2D**). Effects were strongly correlated (Spearman rho=0.65), illustrating broad concordance between methods, with some exceptions.

**Figure 2.**
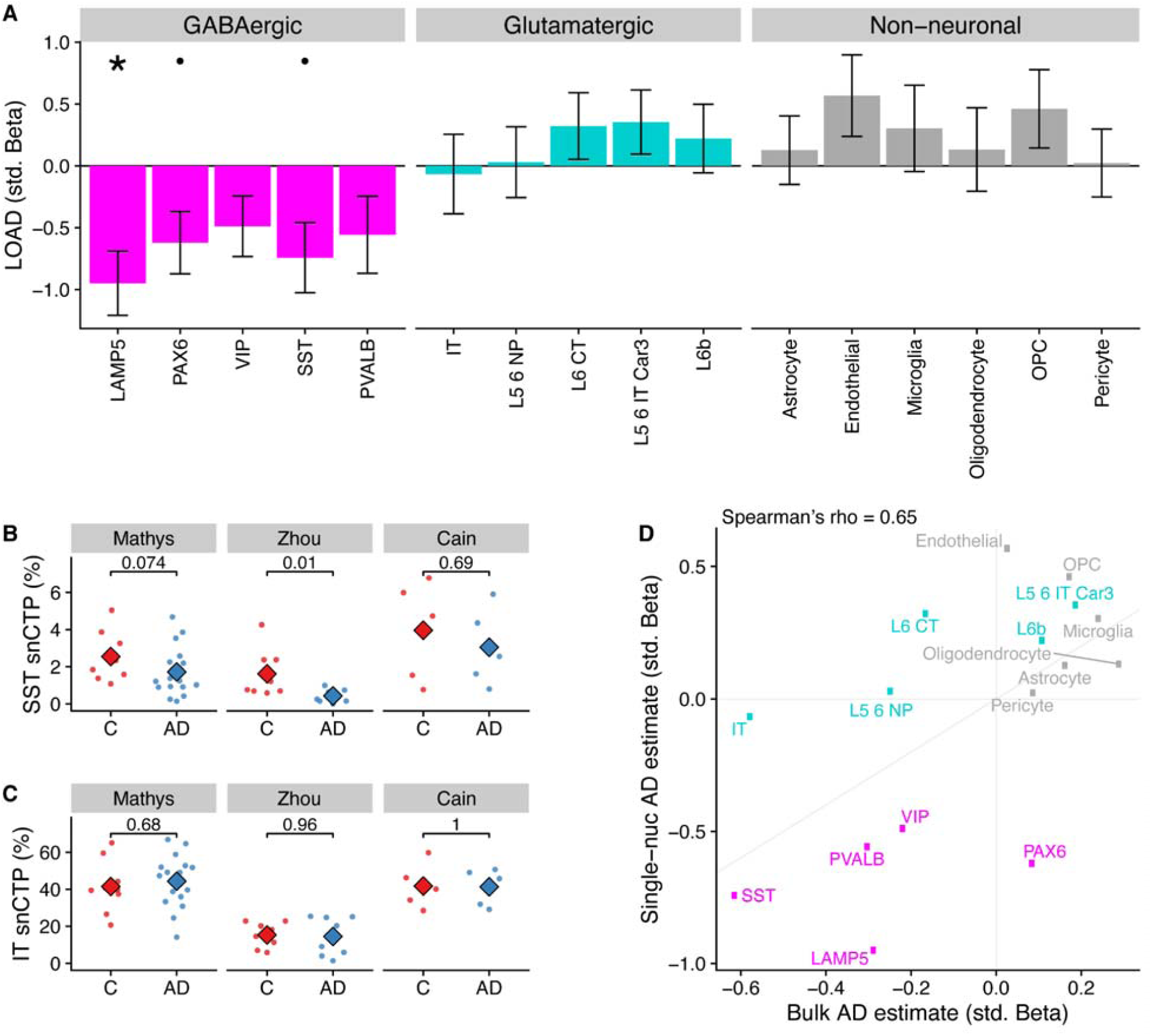
Differences in single-nucleus derived cell type proportions (snCTPs) of neuronal and non-neuronal subclasses in Alzheimer’s disease. A) Consensus associations of AD status and snCTPs across three AD snRNAseq case/control datasets. Y-axis shows standardized beta coefficients estimated using a mixed effects model to pool associations across datasets. Positive (negative) standardized beta coefficients indicate an increase (decrease) in the cell type-specific snCTP in AD. Error bars indicate one standard deviation. B) Comparisons between snCTPs between controls and AD cases in each of three snRNAseq datasets. Y-axis dots show snCTPs for somatostatin (SST) interneurons (as a percentage of all nuclei sampled) from individual post-mortem samples. Numbers show p-values from t- test (uncorrected for multiple comparisons across datasets and cell types) from a statistical model comparing snCTPs between controls and AD cases. Subjects with outlier values of rCTPs not shown. C) Same as B for intratelencephalic-projecting (IT) excitatory pyramidal cells (IT cells). D) Consistency of AD standardized effect sizes between bulk rCTPs and snCTPs based on single-nucleus analyses. X-axis shows point estimates of standardized beta coefficients of AD effects on rCTPs in the ROS/MAP cohort (as in **Figure 1A**) and y-axis is the same as point estimates shown in A. Diagonal line denotes the unity line. Inset correlation value denotes overall Spearman correlation (rho) between rCTP and snCTP estimated effects.

### SST interneurons and IT-projecting pyramidal cells specifically are correlated with AD neuropathologies and residual cognition

Having identified SST interneurons and possibly IT-projecting pyramidal cells as especially diminished in AD cases vs. controls, we explored the associations of rCTPs with specific aging-related neuropathologies and rates of longitudinal cognitive decline. We utilized a larger set of individuals from ROS/MAP with available data (889 subjects), as opposed to only those meeting the consensus AD case/control criteria. After FDR correction, we observed a striking pattern of association whereby only SST rCTPs were negatively associated with each tau and beta-amyloid-related neuropathology - most strongly with neuritic plaques (*p*_FDR_=3.1×10^−4^) - and positively associated with rates of cognitive decline (*p*_FDR_=3.9×10^−6^) and cognition measured proximal to death (*p*_FDR_=5.7×10^−5^) (**Figure 3**). In contrast, IT neurons were also associated with both cognitive measures (decline: *p*_FDR_=8.3×10^−5^; proximal to death: *p*_FDR_=1.2×10^−7^) but were not associated with canonical AD-related neuropathologies. However, an association with TDP-43 proteinopathy was observed (*p*_FDR_=0.015). At a relaxed FDR<0.1 threshold, several other neuronal and non-neuronal associations were observed for individual pathologies (**Figure 3**), though none as strong as those for SST and IT neurons.

**Figure 3:**
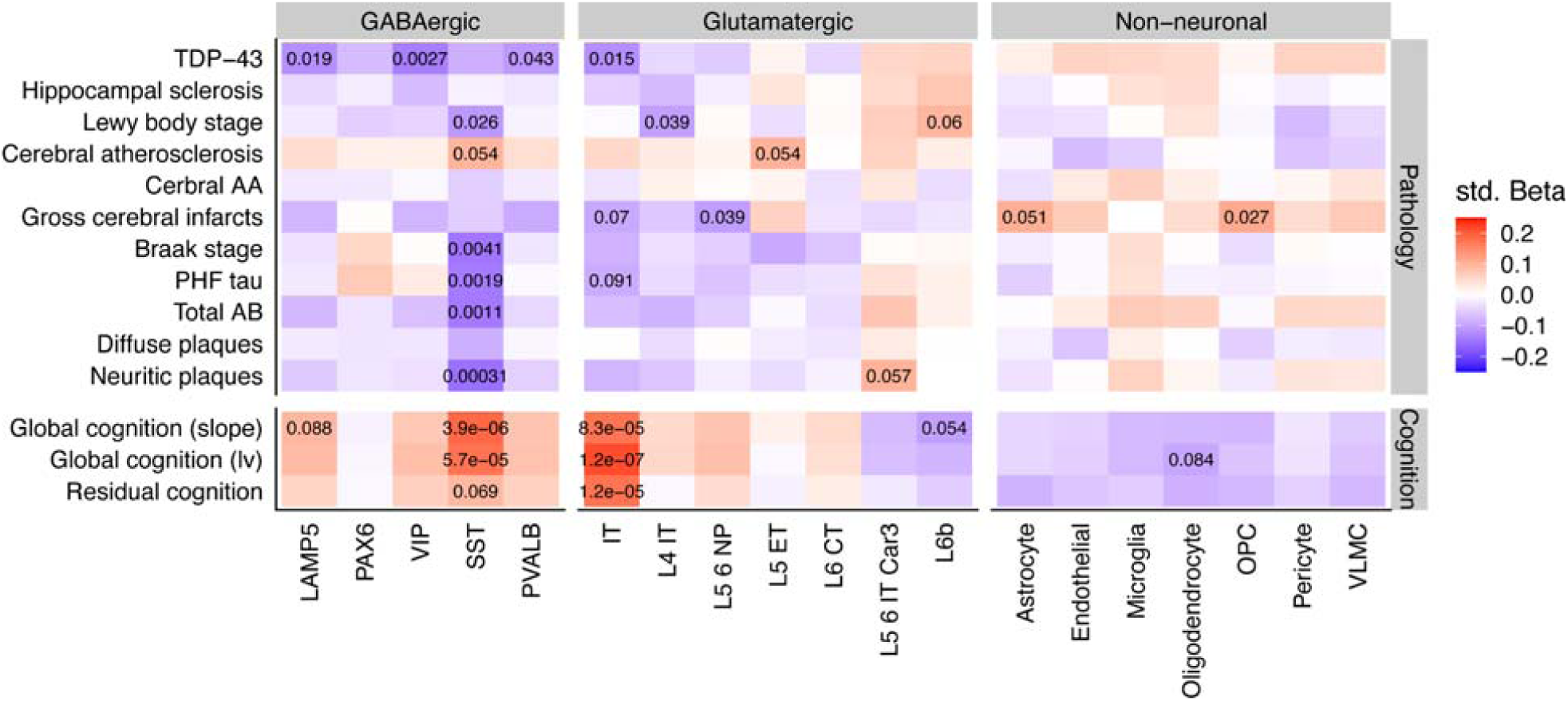
Associations between cell type specific relative proportions and neuropathologies, cognition, and residual cognition. Inset values denote the FDR statistics of specific associations, where FDR < 0.1. Note that while pathology scores are coded such that greater levels of pathology indicate worsening brain health, global cognition scores are coded such that higher scores indicate better cognition and less dementia. Std Beta = standardized beta coefficients.

Finally, we sought to determine if cell type-cognition associations were either independent or a reflection of accumulating brain pathology. Therefore, we calculated a measure of residual cognition for all individuals, as described previously [27], which represents global cognitive performance (proximal to death) after accounting for variability due to neuropathology and demographics (see Methods). After correction, IT rCTPs were the only cell type significantly associated with residual cognition (*p*_FDR_=1.2×10^−5^), though we note SST neurons showed a marginal association as well (*p*_FDR_=0.069). To quantify the additional variance in cognition explained by IT rCTPs over and above measured neuropathology, we first established a baseline model of cognition, where demographic and neuropathological variables alone explained 40.3% of total variance (R^2^=0.403). Adding IT rCTPs to this model increased the variance explained by 1.9% (likelihood ratio test *p*=7.1×10^−8^, R^2^=0.422). By contrast, SST rCTPs increased the explained variance to a much lesser extent (additional 0.53%; likelihood ratio test *p*=0.0049). These findings were supported by mediation analyses, including *APOE* ε4 AD risk variant as a co-variate, which found bidirectional mediation of SST and AD pathology on cognitive performance proximal to death (*p*_perm_<0.0001), but no mediation of the relationship between pathology and cognition by IT neurons (*p*_perm_=0.31), or of IT neurons and cognition by pathology (*p*_perm_=0.35) (results in **Supplementary Table 2**).

## Discussion

Our analysis leveraged three aging and AD studies with multi-region post-mortem bulk gene expression to determine which neocortical cell subtypes are most strongly associated with AD. Based on marker gene expression specific to cellular subclasses, we observed lower relative proportions of most neuronal types and greater relative proportions of most non-neuronal types. In particular, our analyses highlighted fewer somatostatin-expressing (SST) interneurons and intra-telencephalic projecting (IT) pyramidal cells in AD that were replicated across studies and neocortical regions. Cellular proportions directly derived from three additional AD case/control single-nucleus RNAseq datasets provided partial corroboration of our bulk-tissue based results, suggesting that such cellular changes are likely the result of cellular loss as opposed to a coordinated global change in cellular identity. The results of our analyses support previous literature implicating the loss of SST interneurons in AD and further indicate that increased proportions of IT pyramidal cells can contribute to cognitive resilience despite the presence of AD neuropathologies.

Our conclusions implicating SST interneurons, a key subpopulation of cortical GABAergic cells that provide synaptic inhibition to pyramidal cells by targeting their distal dendrites [33], are consistent with a broad literature on the role of SST in neurological illness, recently reviewed [34]. The association of SST with AD is decades-old, beginning with seminal findings reporting reduced levels of somatostatin immunoreactivity in AD brain [35], and more recently with cross-study differential expression analyses finding ubiquitous reductions of SST RNA in AD brain tissue (with the exception of the cerebellum; https://agora.adknowledgeportal.org/). However, it remains unknown if this association is driven by a selective loss of SST mRNA or losses of populations of SST-expressing neurons. Taken as a whole, our bulk and single-nucleus based findings support the latter conclusion, though certainly do not provide definitive evidence for it. In agreement, one study found that SST interneurons were uniquely lost in AD among neuronal subtypes in prefrontal cortex ROS/MAP samples [12]. While the precise role of selective SST interneuron vulnerability in AD remains to be understood, a recent publication pointed to a role for a potential direct biochemical interaction between the SST neuropeptide and amyloid beta [36].

We also observed strong associations between IT pyramidal cells and AD, and, intriguingly, this was the only cell type significantly associated with residual cognition. IT cells are defined by their cortico-cortical and cortico-striatal projections [31] and encompass supragranular pyramidal cells, such as CUX2-positive cells, and infragranular cells, including RORB- and THEMIS-positive pyramidal cells [24]. Immunohistochemical studies corroborate these results in part, suggesting that SMI32-immunoreactive neurons, labelling large pyramidal neurons in Layers 3 and 5, are selectively lost in the frontotemporal cortex in AD [37–39]. More recently, evidence from snRNAseq studies of the entorhinal cortex and superior frontal gyrus have implicated RORB-expressing excitatory neurons as selectively vulnerable in AD [15]. As one caveat, we note that we saw some, but overall limited replicability of decreased IT CTPs in our bulk-compared to our single-nucleus analyses.

Our study has several key limitations and considerations. First, the backbone of our study is a neocortical cell type taxonomy derived from deep transcriptomic sequencing of single-nuclei from normotypic individuals [24]; it remains unclear how comparable these transcriptional profiles are to those in the elderly and in individuals with AD. Second, the conclusions of our study rely on the accurate estimation of relative differences in cell-type proportions between individuals. Such estimates are highly dependent on the choice and quality of the constituent marker genes that serve as representatives of each cell type [4,5] as well as the particular choice of method for cellular deconvolution [8,9,32]. However, the relatively high degree of consistency between our bulk- and single-nucleus based results help partially mitigate this concern. Third, the focus of this work is the study of robust changes in cell-type proportions in AD, but does not tackle the question of within-cell type transcriptional regulation [13,40]. Lastly, all of the results presented here are associational; further studies are needed to determine how and when cell type-specific loss occurs relative to the emergence of AD pathologies and cognitive decline.

## Conclusions

Overall, our study provides a comprehensive consensus overview of the vulnerability of neocortical neuronal subpopulations in AD. Our results demonstrate that losses of SST interneurons and IT pyramidal cell populations are those most strongly associated with AD. In addition, IT pyramidal cells are uniquely associated with residual cognition, suggesting that efforts to preserve or maintain this key neuronal subpopulation might mitigate cognitive decline in the face of AD neuropathologies. Our hope is that these results will inform future studies to further disentangle the causal progression of AD neuropathological burden, cellular loss, and cognitive decline.

## List of Abbreviations

AD: late-onset Alzheimer’s disease
CERAD: Consortium to Establish a Registry for Alzheimer’s Disease
FDR: false discovery rate
GABA: gamma aminobutyric acid
rCTP: relative cell-type proportion
snCTP: single-nucleus cell type proportion
IT: intratelencephanic
MAP: Rush Memory and Aging Project
MSBB: Mount Sinai Brain Bank
ROS: Religious Orders Study
SST: somatostatin
TMM: trimmed mean of m-values

## Declarations

### Ethics approval and consent to participate

For The Religious Orders Study and Rush Memory and Aging Project, all study participants provided informed consent and both studies were approved by a Rush University Institutional Review Board. Further, all participants signed an Anatomic Gift Act for organ donation and signed a repository consent for resource sharing. For the Mayo dataset, protocols were approved by the Mayo Clinic Institutional Review Board and all subjects or next of kin provided informed consent.

### Consent for publication

Not applicable.

### Availability of data and materials

The RNAseq datasets supporting the conclusions of this article are available via approved access at the Synapse AMP-AD Knowledge Portal (https://adknowledgeportal.synapse.org/, doi: 10.7303/syn2580853). All analyses were performed using open-source software. No custom algorithms or software were used that are central to the research or not yet described in published literature. ROSMAP resources can be requested at https://www.radc.rush.edu.

### Competing interests

The authors declare no conflicts of interest. Funders did not play any role in the design, analysis, or writing or this study.

### Funding

Funding support for DF was provided by The Koerner Family Foundation New Scientist Program, The Krembil Foundation, the Canadian Institutes of Health Research, and the CAMH Discovery Fund. MEC, YC, and SJT acknowledges the generous support from the CAMH Discovery Fund, Krembil Foundation, Kavli Foundation, McLaughlin Foundation, Natural Sciences and Engineering Research Council of Canada (RGPIN-2020-05834 and DGECR-2020-00048), and Canadian Institutes of Health Research (NGN-171423 and PJT-175254), and the Simons Foundation for Autism Research. ROSMAP was supported by NIH grants P30AG10161, P30AG72975, R01AG15819, R01AG17917, U01AG46152 and U01AG61356.

### Authors’ contributions

MC was responsible for data processing, statistical analysis, manuscript writing, and editing. YC contributed to statistical analyses, data visualization, and manuscript writing. DF and SJT were responsible for data access, ensuring data quality control, study design, and manuscript writing and editing. VM, YW, PLDJ, DAB, and JAS were responsible for aspects of data collection, collaborative input on study design, and manuscript editing.

## Acknowledgements

The authors acknowledge all of the patients and their families for graciously donating brain tissue. We thank Lilah Toker and members of the Felsky and Tripathy laboratories for comments on this work and Milos Milic for assistance with data management.

## Notes

### Competing Interest Statement

The authors have declared no competing interest.

