## Supplementary Information for "Bulk and Single-nucleus Transcriptomics Highlight Intra-telencephalic and Somatostatin Neurons in Alzheimer’s Disease"

#### Supplementary Methods

##### Bulk RNA sequencing in ROS/MAP

All samples were extracted using Qiagen's miRNeasy mini kit (cat. no. 217004) and the RNase free DNase Set (cat. no. 79254), and quality was evaluated by Agilent Bioanalyzer. Briefly, for pipeline #1, The Broad Institutes's Genomics Platform performed RNA-Seq library preparation using the strand specific dUTP method30 with poly-A selection31. Sequencing was performed on the Illumina HiSeq with 101bp paired-end reads and achieved coverage of 150M reads of the first 12 samples. The remaining samples were sequenced with coverage of 50M reads. For pipeline #2, RNA sequencing libraries were prepared using the KAPA Stranded RNA-Seq Kit with RiboErase (kapabiosystems) in accordance with the manufacturer’s instructions. Sequencing was performed on the Illumina NovaSeq6000 using 2 x 100bp cycles targeting 30 million reads per sample. For pipeline #3, RNA was extracted using Chemagic RNA tissue kits (Perkin Elmer, CMG-1212) on a Chemagic 360 instrument. 500ng total RNA was used as input for sequencing library generation and rRNA was depleted with RiboGold (Illumina, 20020599). A Zephyr G3 NGS workstation (Perkin Elmer) was utilized to generate TruSeq stranded sequencing libraries (Illumina, 20020599). Sequencing was performed on a NovaSeq6000 (Illumina) at 40-50M reads (2x150bp paired end). Full details on these methods are available on the AMP-AD knowledge portal (syn3219045).

##### Single-nucleus RNAseq datasets

In total, we accessed expression data from four human cortical single-nucleus RNAsequencing (snRNAseq) datasets for this study.

For cell type marker gene development, ultra high-depth single-nucleus RNA sequencing data from the human neocortex was obtained from datasets provided by the Allen Institute for Brain Sciences (AIBS). Specifically, we made use of the “Multiple Cortical Areas - Smart-seq (2019)” dataset accessed in August 2020 (<https://portal.brain-map.org/atlases-and-data/rnaseq/human-multiple-cortical-areas-smart-seq>) with collection and analysis methodology described in Hodge et al. 2019 [[1]](https://www.zotero.org/google-docs/?zVRgnO). Briefly, single nuclei were sorted and reverse transcribed using SMART-Seq.v4 protocol. Outlier cells were removed according to the “outlier_call” column in accompanying metadata. For our analyses, we used all nuclei sampled from the cingulate gyrus (5,939 nuclei) and medial temporal cortex (15,519), as these correspond most closely with the bulk expression samples described above. Given that nuclei from non-neuronal cell types were relatively undersampled in these datasets (722 from cingulate gyrus and 1,411 from medial temporal cortex), we supplemented with 2,620 nuclei corresponding to non-neurons sampled from other cortical regions, including visual, auditory, somatosensory and motor cortex (502, 742, 595, and 781 nuclei respectively).

For analyses of snRNAseq-derived cell type proportions in AD, three snRNAseq datasets collected from individual subjects sampled from the ROS/MAP cohort were also used.

1. Mathys et al [[2]](https://www.zotero.org/google-docs/?RvWZrN), sampled from 48 individuals. Processed data were obtained from Synapse (ID: syn18681734). Specifically, we made use of files labeled “filtered_count_matrix.mtx” and “filtered_column_metadata.txt”, which reflected 70634 cells passing quality control (described in [[2]](https://www.zotero.org/google-docs/?xtX697)).
2. Zhou et al [[3]](https://www.zotero.org/google-docs/?QA8NCJ), reflecting 32 individual subjects. We processed and analyzed data from 32 10X-genomics-format matrices with their accompanying metadata (“snRNAseqAD_TREM2_assay_scRNAseq_metadata.csv” and “snRNAseqAD_TREM2_biospecimen_metadata.csv”), also from synapse.org (ID: syn21670836). As the cells from these files were not pre-filtered based on quality control metrics, we performed a simple quality-control step by selecting cells that expressed 200-2500 genes after mapping cells to the AIBS cell taxonomy as described below. After filtering, this Zhou et al dataset consisted of 89993 cells. The authors state that their RNAseq data were aligned to the “hg38, human genome GRCh38” reference.
3. Cain et al [[4]](https://www.zotero.org/google-docs/?RXNVuM), reflecting 24 individual subjects. Processed data were obtained from “ROSMAP_Brain.snRNAseq_metadata_cells_20201107.csv", “ROSMAP_Brain.snRNAseq_metadata_genes_20201107.csv”, and “ROSMAP_Brain.snRNAseq_counts_sparse_format_20201107.csv.gz” (synapse ID: syn16780177). The dataset contains 162767 cells. The authors state that their RNAseq data were aligned to “the hg38 transcriptome”. Cells that the authors labelled as "None.NA" for their cell-type identity were considered low quality and removed from further analysis.

##### Description of Cell Type Taxonomy

We made use of the cortical cell type taxonomy recently established by the Allen Institute for Brain Sciences (AIBS) that defines taxonomic relationships between cells and cell types in the human neocortex [[1]](https://www.zotero.org/google-docs/?LQgxLm). This taxonomy was based on unbiased clustering analysis of snRNAseq datasets and uses consistent nomenclature with orthologous datasets from the mouse neocortex, enabling annotation of snRNAseq derived clusters with multi-modal features and attributes. We focused our analyses primarily on the “subclass” cell type resolution (e.g., somatostatin-expressing GABAergic interneurons), which serves as an intermediate resolution between more coarse-grained (e.g., excitatory versus inhibitory neurons) and fine-grained cell taxonomic divisions (e.g., Martinotti neurons). Taxonomy information can be found here: https://portal.brain-map.org/atlases-and-data/rnaseq/human-multiple-cortical-areas-smart-seq.

Cell types were primarily operationally defined at the “subclass” resolution as defined by AIBS, using the column “subclass_label”. We similarly made use of cell class definitions provided by AIBS via the “class_label” column, which identified cells as “GABAergic”, “Glutamatergic”, and “Non-neuronal”. Finally, we also combined the GABAergic and Glutamatergic cells defined by AIBS class labels into a “Neuronal” group. We used AIBS defined subclasses, classes, and our AIBS-derived “Neuronal” group for the marker selection and/or subsequent analyses described below.

##### Mapping of external snRNAseq data onto AIBS cell taxonomy

Cells from each of the three ROS/MAP snRNAseq datasets (Mathys, Zhou, and Cain) were mapped onto the AIBS snRNAseq dataset after the cell selection described under “Single-cell RNAseq dataset description”, including filtering out low-quality outlier cells. Cross-dataset mapping was performed using the FindTransferAnchors and TransferData functions in Seurat version 3.2.1 after identification of the top 20 percent of most variable features (genes) for each dataset [[5]](https://www.zotero.org/google-docs/?IzPjrq). snRNAseq nuclei were mapped to subclass labels from the AIBS reference dataset. Individual nuclei with mapped cell type identities with Seurat-derived prediction scores greater than 0.5 were accepted, except for nuclei mapping to the AIBS “IT” subclass, where a more stringent cutoff of 0.8 was used. We found this additional stringent cutoff necessary, because in preliminary analyses, nuclei mapping to the IT cell group had more unexpected mappings to mis-matched cell types (compared to original identities in Cain et al. and Mathys et al.) than other reference subclasses. Following this custom re-mapping, we retained 54880 nuclei from Mathys, 50417 nuclei from Zhou, and 128094 nuclei from Cain.

Confusion matrices were used to quantify the overlap between original-author annotated cell type labels and our re-mapped labels following mapping to the AIBS reference taxonomy. We note that original-author annotated cell type labels were not provided for the Zhou et al dataset.

##### Generation of cell type-enriched marker genes

Cell-type marker identification from snRNAseq data was performed similarly to previously published methods [[6]](https://www.zotero.org/google-docs/?jXgSIg). Marker genes for the taxonomic groupings specified previously (AIBS-defined cell subclasses and classes and the AIBS-derived “Neuronal” group) were identified by differential expression testing between taxonomic groups of the same resolution, such as “SST” cells compared to cells of all other subclasses or “GABAergic” cells vs “Glutamatergic” and “Non-neuronal” cells. “Neuronal” cells were also compared to “Non-neuronal” cells to transcriptomically define markers of neurons. Differential gene expression tests were conducted using Seurat Version 3.2.1 [[5]](https://www.zotero.org/google-docs/?t9bTxn) in R version 3.6 (R Core Team, 2019). Specifically, marker genes were identified using the FindAllMarkers function with a log fold-change threshold of 2.5 and a minimal detection percentage of 35% (min.pct = 0.35). Both the “MAST” and “roc” methods were used with log-normalized gene counts, and a marker detected with either method was included in the potential marker list. In cases where potential markers were shared between multiple cell types, such genes were removed as markers from our analysis to yield a final marker list with high specificity.

##### Estimation of relative cell type proportions from bulk RNAseq samples

Relative cell type proportions were estimated with the MarkerGeneProfile (MGP) R package, as described previously [[7,8]](https://www.zotero.org/google-docs/?Bf9IOd), using our AIBS-derived human neocortex cell type-specific marker genes described above. The output of the mgpEstimate function was taken as the relative cell-type proportion estimates (rCTPs), providing an indirect measure of cell type abundance in each sample. To ensure consistency in rCTP definitions across individual bulk datasets, rCTPs were estimated using only marker genes passing QC in all datasets. rCTPs were converted to standardized z-scores within each dataset prior to downstream analysis.

##### Estimation of RNA seq-derived cell type proportions

Cell type proportions from snRNAseq datasets (snCTPs) were directly estimated from snRNAseq datasets by counting nuclei annotated to each cell type and normalizing by the total count of all QC-passing nuclei per individual subject. We note that such calculations were only performed on nuclei passing quality control and also met our mapping criteria to our reference cell type taxonomy. Direct comparisons between bulk and snRNAseq derived cell type proportions for subjects from the ROS/MAP cohort were performed by identifying subjects in common between both datasets and correlating rCTPs with snCTPs values across subjects.

##### Differential gene expression using bulk tissue data

Association analysis of bulk tissue gene expression and AD case/control status were performed as described previously [[9,10]](https://www.zotero.org/google-docs/?c7z54s). This was done using limmas lmFit, modelMatrix, and eBayes methods in R. Before running removeBatchEffect and after calculation of TMM normalization values and mean-variance derived observational-level weights, each dataset (ROS/MAP, Mayo, MSBB FP, MSBB STG, MSBB PHG, and MSBB IFG) was inputted to lmFit for a robust regression with maximum number of iterations capped at 1000. The design matrix for the lmFit method was constructed using limma’s modelMatrix function, accounting for sequencing batch, sex, percent of mapped bases, percent usable bases, RNA integrity number (RIN), age at death, postmortem interval and AD diagnosis. Limma’s eBayes method was then used on the result of the lmFit call for each dataset to calculate moderated t-statistics, moderated F-statistics, and log-odds of differential expression. From the resulting object the strength (p-value) of the association between each gene present in the dataset and AD diagnosis was extracted. The beta-coefficient, or direction of the association between each gene in the dataset and AD diagnosis, was derived from the object returned by the lmFit call.

#### The genes with p-values and beta-values calculated from their association with AD diagnosis were then subset into marker gene categories for each cell-type defined by the human neocortex snRNAseq-derived cell type-specific marker genes. Each marker gene’s strength and direction of association to AD diagnosis could then be determined per cell-type, per dataset.

##### Differential gene expression using single-nucleus data

To estimate differences in average SST mRNA expression per SST-annotated nucleus between AD cases and controls, we first calculated the average normalized expression of SST mRNA among each SST-annotated nucleus per subject. Only subjects with at least 3 QC-passing SST nuclei were included in analysis.

We used a linear model to estimate the effect of AD diagnosis on normalized SST expression (among SST-annotated nuclei) while covarying for each subject’s SST cell type proportion and count of SST-annotated nuclei, in addition to snRNAseq dataset identity, age at death, sex, and post-mortem interval. For plotting purposes, we residualized SST mRNA expression using a model identical to the one below, but excluding the term for AD diagnosis.

(A) SST_mRNA_expr ~ AD diagnosis + SST_cell_proportion + SST_cell_count + dataset +

age at death + sex + pmi

#### Supplementary Figures

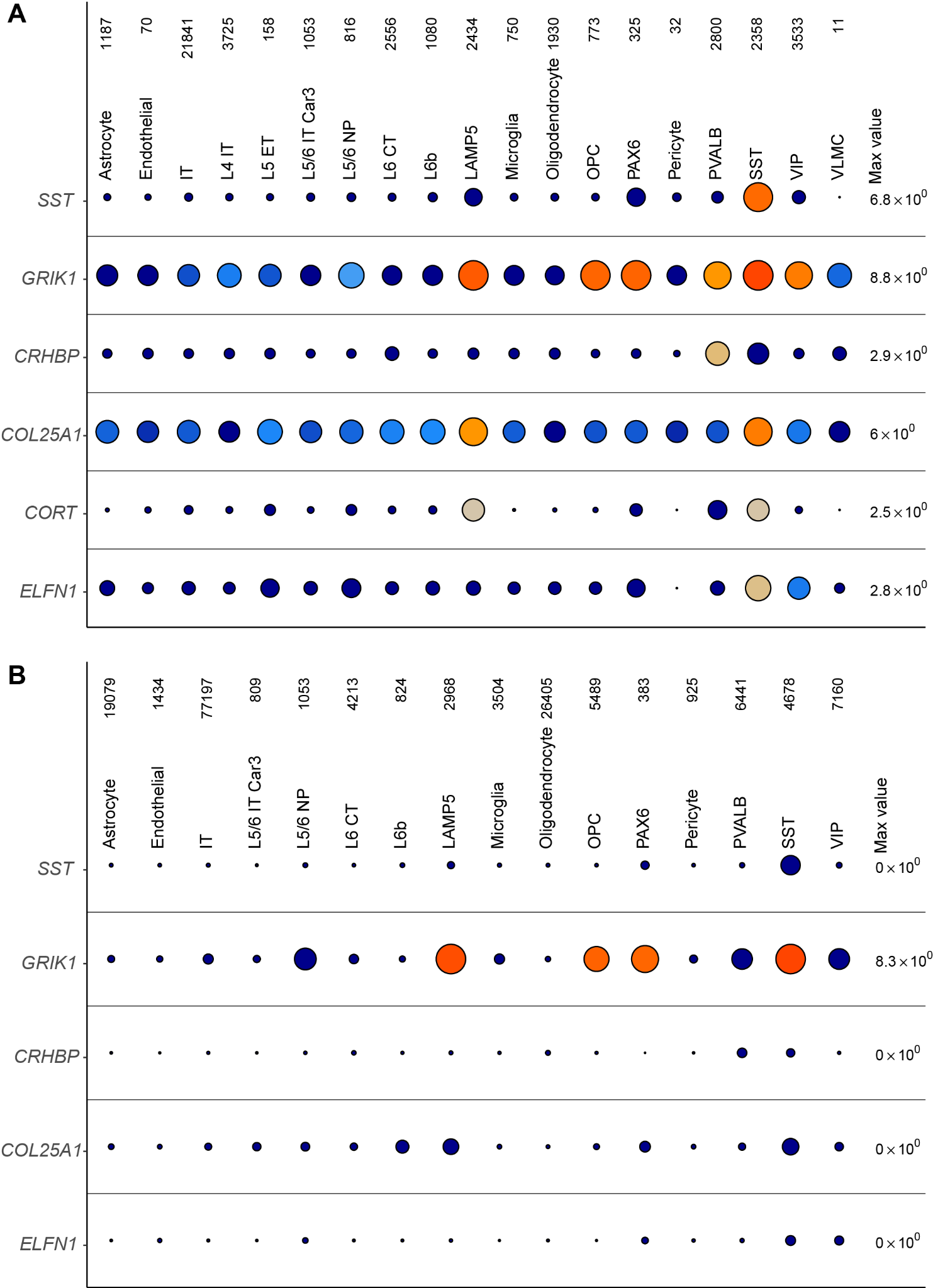

*Supplemental Figure 1: Group dot plots showing consistency and replicability of gene expression between Allen Institute snRNAseq dataset (A) and Cain single-nucleus AD dataset (B) for SST cells.*

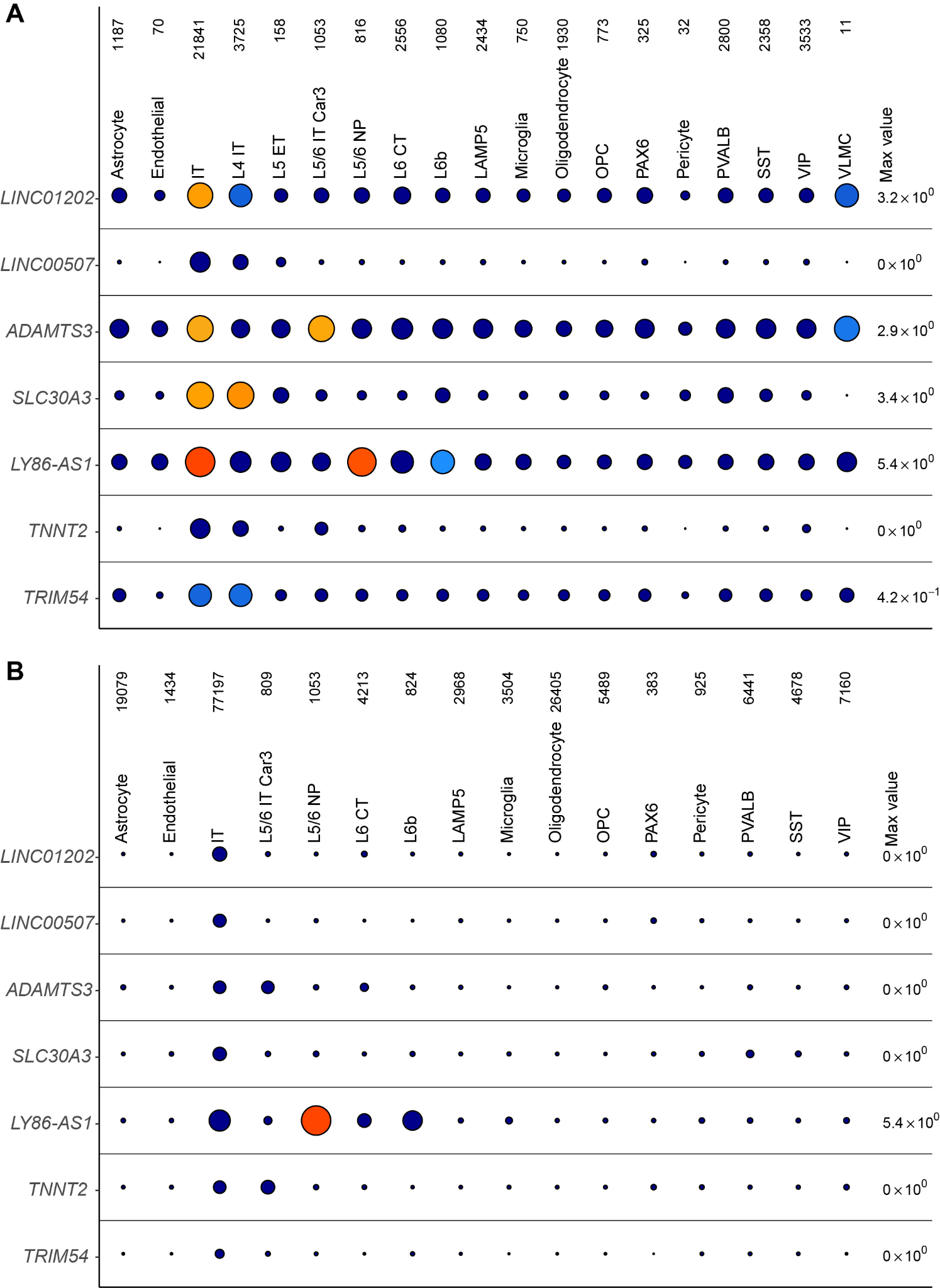

*Supplemental Figure 2: Group dot plots showing consistency and replicability of gene expression between Allen Institute snRNAseq dataset (A) and Cain single-nucleus AD dataset (B) for IT cells.*

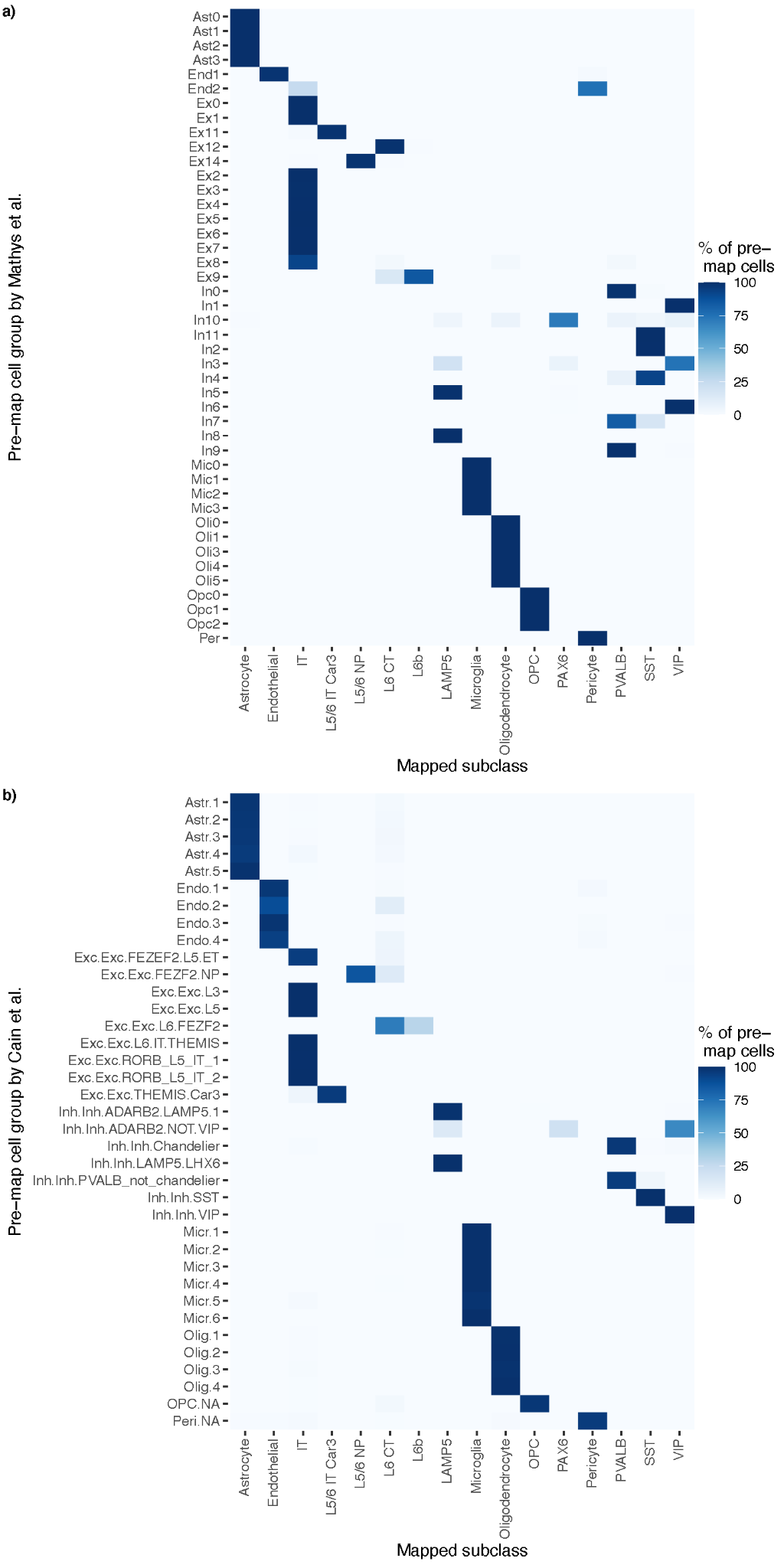

*Supplemental Figure 3: A) Comparison of cell type identities based on annotations of single-nucleus RNAseq data from Mathys et al (A) and Cain et al (B). Rows show original cell type annotations from Mathys and Cain and columns show re-mapped cell identities to the Allen Institute reference cell type taxonomy used in this publication. Rows sum to 100%. Note that only single-nuclei which are re-mapped to the reference taxonomy following quality filtering are shown.*

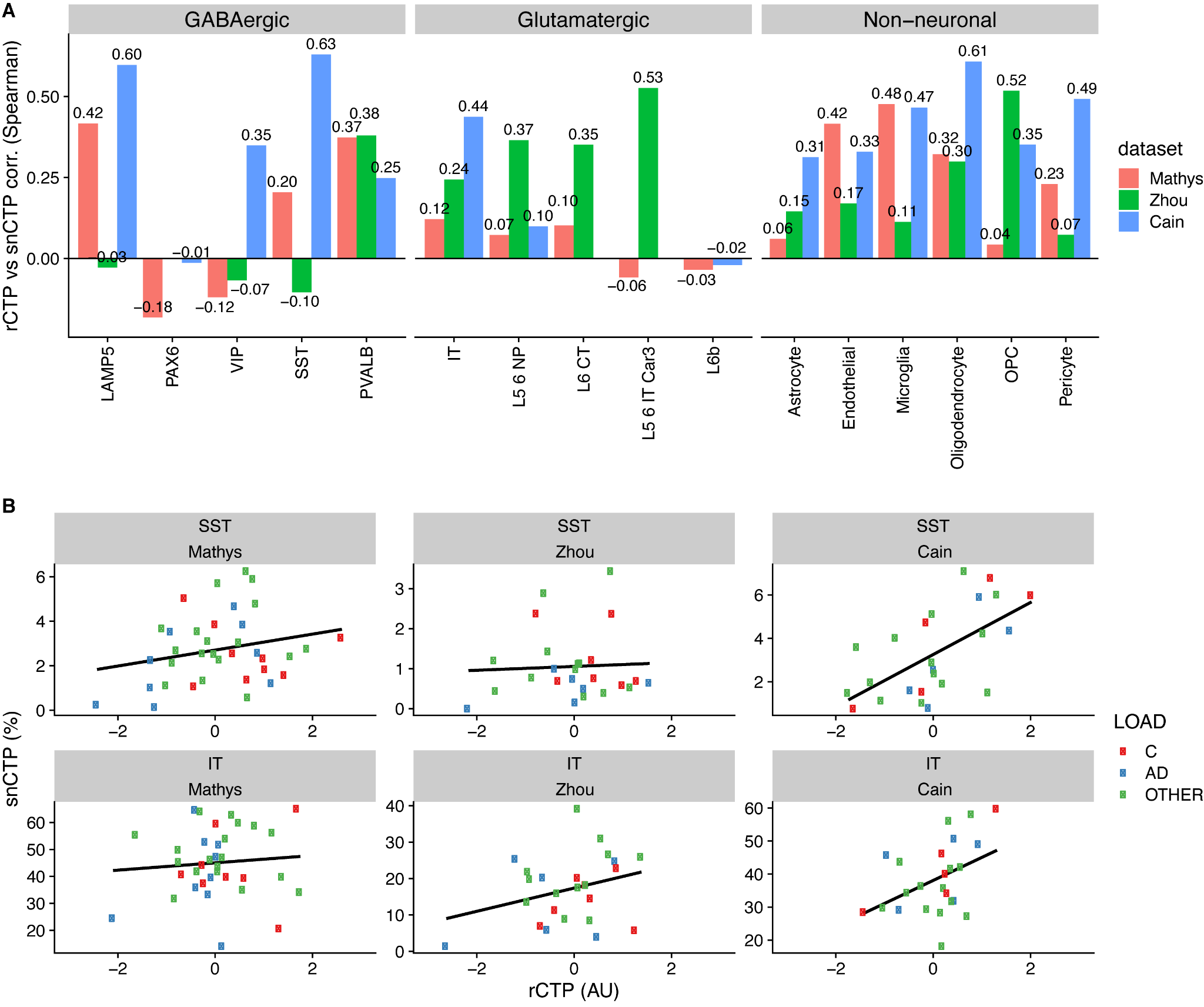

*Supplemental Figure 4. Consistency between bulk and single-nucleus derived cell type proportions. A) Summary of Spearman correlations between rCTP and snCTPs based on subjects shared between analyses bulk and snRNAseq analyses. Correlations calculated independently using subjects shared between Mathys, Zhou, and Cain snRNAseq datasets. B) Scatterplots illustrating consistency between snCTPs (y-axis, with cell type proportions calculated as percentages) and rCTPs (x-axis, arbitrary units) for SST cells and IT cells. For SST interneurons, the correlations between rCTPs and snCTPs derived from the Mathys, Zhou and Cain datasets were Spearman’s rho = 0.20, -0.10, and 0.63 respectively. Similarly, for IT glutamatergic cells the correlation between rCTPs and snCTPs derived from the Mathys, Zhou and Cain datasets were Spearman’s rho = 0.12, 0.24, and 0.44 respectively. These analyses suggest broad comparability between both methods (bulk and snRNAseq-based) for estimating cellular proportions as well as key differences between the different single-nucleus RNAseq datasets that might influence our snCTP calculations.*

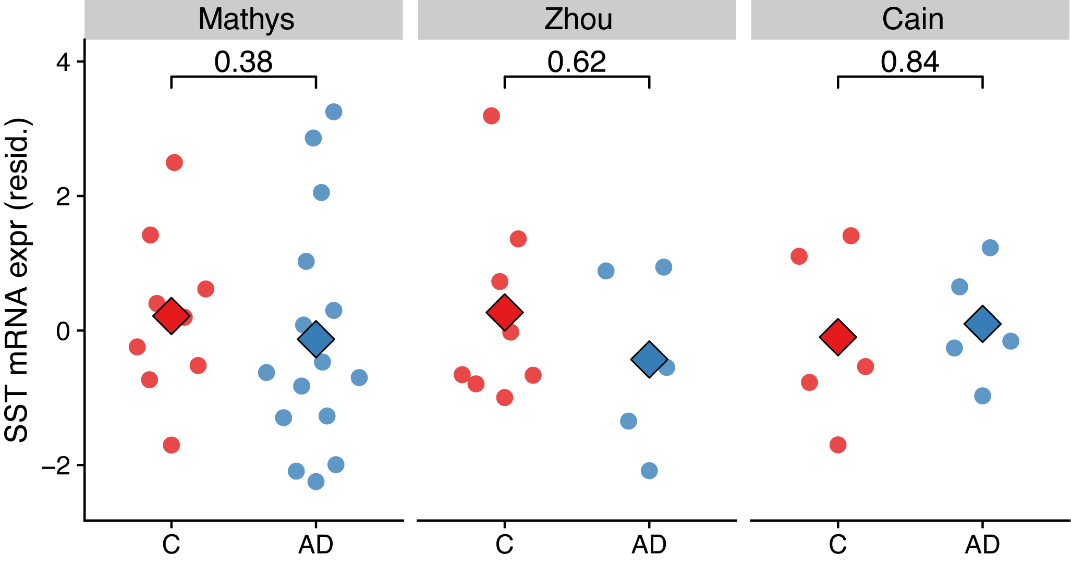

### *Supplemental Figure 5. SST mRNA expression is not significantly different between SST cells from AD cases versus controls. Y-axis shows mean SST mRNA expression per nucleus among nuclei that were bioinformatically annotated to the SST cell type. Y-axis shows residualized expression values after covarying, per subject, for age at death, sex, and post-mortem interval as well as count of SST positive nuclei SST single-nucleus proportion. Each dot reflects one subject and large diamonds indicate group means. Only subjects with at least 3 QC-passing SST nuclei were included in analysis. Inset values indicate uncorrected p-values based on a Wilcoxon rank sum test. Mega-analysis for AD diagnosis term (encompassing across all three datasets): std. Beta = -0.428, pval = 0.417, n = 47 subjects.*

#### Supplementary Tables

<https://github.com/sonnyc247/MarkerSelection/blob/master/Data/Outputs/CSVs_and_Tables/Markers/MTG_and_CgG_lfct2/new_MTGnCgG_lfct2.5_Publication.csv>

*Supplementary Table 1*. *List of cell type-specific markers used in estimating relative cell type proportions.*

*Supplementary Table 2.* *Results from mediation modelling of global cognition, neuropathology, and SST and IT neurons proportions in ROS/MAP. Note: ci = 95% confidence interval; cogn_global_lv = global cognitive performance measured proximal to death; gpath_sqrt = sqare-root transformed global AD neuropathology (see Methods); propmed = proportion of total effect mediated.*
